# A Pilot Study on the Urinary Microbiome Composition and Diversity in Clinical UTI Samples: A 16S rRNA Analysis

**DOI:** 10.64898/2026.04.18.719336

**Authors:** Musaab Hamid Farhan, Ayad M.J. Al-Mamoori, Noor S.K. Al-Khafaji, Aws H. J. Al_Rahhal

## Abstract

This pilot study assessed the composition and diversity of the urinary microbiome from clinically confirmed UTI samples using 16S rRNA sequencing, whilst also exploring inter-individual variability of microbial community structure.

We examined ten urine samples from patients with culture-positive UTIs. Demographic and clinical metadata, including age, sex, body mass index (BMI), diabetes status and recent antibiotic exposure was recorded per sample. Metagenomic DNA was extracted from microbial samples and sequenced to generate genus-level taxonomic profiling through 16S rRNA gene sequencing. Relative abundance tables were generated for each of the samples to identify dominant bacterial genera within each sample and summarize cohort level microbial patterns. To evaluate within-sample richness and evenness, alpha diversity indices (Shannon, Simpson, observed features and Chao1) were computed; beta diversity was measured using Bray–Curtis dissimilarity with principal coordinates analysis (PCoA) for graphical representation.

The study’s findings revealed the sex and moderate clinical diversity of the study sample; all samples were confirmed as having been taken from a UTI patient and exhibited a wide level of heterogeneity regarding the microbial composition of each urine sample. Overall, Pseudomonas was the dominant genus present, however, specific samples had approximately 50% of their microbiomes composed of Klebsiella, Proteus, and Escherichia species as well as approximately 25% of their total microbes were made up of Burkholderia spp., which are closely related to the genus of interest used during the course of this study. The observed alpha diversity of each sample displayed considerable variation for the included samples with a continuum of samples ranging from a single dominant microbe to a highly diverse mixed population producing a highly diverse polymicrobial population/bacterial composition, with some ratios of individual taxa to collective taxa of many samples repeatedly illustrating the exact nature of the specimen. Furthermore, a significant degree of Beta diversity was found between the patients, providing compelling evidence of identifiable differences among urinary microbiomes between patients with UTI.

This pilot project provides a clear indication of the diversity and overall heterogeneity of urinary microbiota found in the UTI patients studied. In addition, the results of this study support the notion that the ecological complexities present within a urinary microbiome cannot necessarily be established through conventional culture methods, and that combined with molecular techniques such as 16S rRNA sequencing of bacterial DNA could be used to quantify and characterize the ecologic condition of urinary microbiota separate from the traditional high prevalence of identifiable uropathogens.

## 1. Introduction

Urinary tract infections (UTIs) are among the most common bacterial infections seen in clinical practice, and they are a leading cause of antibiotic use, recurrent morbidity, and healthcare burden around the world. They range from simple lower urinary tract infections to complex and recurrent manifestations that are frequently attributed to underlying host factors, including metabolic disorders, prior antimicrobial exposure, as well as structural or functional abnormalities of the urinary tract(1,2). While urine culture is the standard gold standard method still used for the diagnosis of UTI, traditional approaches based on culture are time consuming and have limited sensitivity for fastidious or slow-growing organisms unable to fully characterize oligobacterial populations(3,4). Such limitations have resulted in growing interest towards molecular approaches that enable a more comprehensive characterization of the urinary microbial community beyond just the dominant culturable pathogen(5,6).

Recent breakthroughs in next-generation sequencing technologies have called into question the traditional notion of the urinary tract as a sterile site, providing evidence for diverse microbial assemblages, termed the urobiome(1,7). Urinary microbiome diversity may underpin urinary tract health and balance, impacting the susceptibility to urinary infection and clinical outcomes(8,9). In particular, the balance between dominant uropathogens and background commensal or opportunistic taxa may dictate the progression of colonization to symptomatic infection among individuals with predisposed clinical conditions(10,11). Thus, accurate high-throughput characterization of urinary microbiome composition has become a promising step in understanding UTI pathophysiology and refining microbial signatures concordant with disease phenotypes(6,12).

16S rRNA gene sequencing allows for culture-independent identification of bacterial genus level and quantification of community structure in each sample(13,14). It yields principal ecological parameters such the relative abundance of dominant genera, the diversity in a sample (alpha diversity), and dissimilarity between samples or groups of samples (beta diversity)(13,15). Taken together, these data provide a fine-grained view of urinary microbial communities and enable investigation of inter-individual variation in UTI-associated microbiota(16,17). The variation could be a consequence of host factor, antimicrobial exposure and metabolic state differences that are risk factors for UTI(18,19).

Here, we describe a pilot analysis of 16S rRNA sequencing-based urine microbiota profiling in samples from subjects prospectively clinically diagnosed with UTI matched against relevant demographics and clinical variables including age, sex, body mass index (BMI), diabetes mellitus diagnosis and results from coinciding urine cultures as well as contemporaneous history of antibiotic exposure. This cohort was well balanced with respect to sex and had a wide age distribution, as well as some moderate metabolic comorbidity. This was especially true as a significant proportion of the study population stated recent exposure to antibiotics a critical driver of microbial diversity and for pathogen domination or dysbiosis in general(20,21). All samples analyzed were derived from culture-positive UTIs, thus ensuring that microbiome findings are directly linked to the clinically defined infectious input and even allow consideration of a wider community than mono-species culture results.

Preliminary microbiome profiling showed evidence of genus-level differences between samples despite a limited number of uropathogenic genera identified as patient-dominant taxa and overall heterogeneity. Some of the samples were almost entirely dominated by a single genus (as you’d expect from classic infections driven by pathogens) but others had much more complex, even mixed microbe assemblages. This heterogeneity underscores the diverse bacterial taxa associated with UTI urine, and the limitation of using a single culture to represent that entire microbial community. Individual microbial genera were over-or underrepresented differentially in groups of samples compared with one another, consistent with recurrent cohort-level summaries that have implied dominant enrichment trends across multiple subjects in UTIs.

Ecologically, alpha diversity analyses showed significant differences in alpha diversity indices between samples, which indicates that urinary microbial communities differ significantly in relation to richness and evenness. Lower diversity profiles were generally found in cases with dominating species of a single genus, while higher diversity profiles suggested more polymicrobial compositions. We observed a marked dissimilarity between patients in our beta diversity analysis, reinforcing the notion that each UTI case may possess its own unique microbial composition shaped by host and clinical variables. Multivariate ordination methods also demonstrated clustering of some samples with comparable community profiles and separation of others, reinforcing the notion of distinct microbiome signatures in UTI.

Here we performed a comprehensive descriptive analysis of urinary microbiome composition and diversity in clinically diagnosed UTI cases using 16S rRNA sequencing, based on these observations. More specifically, we aim to (i) describe baseline demographic and clinical characteristics relevant to urinary microbial diversity; (ii) identify the dominant and prevalent bacterial genera across urine samples; and (iii) assess within-sample diversity of taxa richness/abundance and between-samples dissimilarity/shifts in microbial communities to reflect inter-individual heterogeneity in the urobiome.

## Materials and Methods

### Study Design and Setting

Based on our 16S rRNA gene sequencing effort, this study was performed as a pilot, observational descriptive analysis of the urinary microbiome composition and diversity identified in clinical UTI samples. This focus on culture-confirmed UTI cases allowed the characterization of microbial community structure beyond the boundaries imposed by traditional microbiological diagnostics.

### Study Population and Sample Size

A total of 10 urine samples (S1–S10) were obtained from patient diagnosed with UTI as part of this pilot analysis. Each patient sample was given a unique identifier that mapped the clinical metadata with the sequencing outputs. The small sample size was selected as this allowed for an early exploratory assessment of urinary microbiome variation and methodological feasibility of larger future studies.

### Inclusion and Exclusion Criteria Inclusion Criteria

Patients were included if they met the following criteria:

1. Clinically diagnosed urinary tract infection.
2. Result of positive urine culture indicating the presence of a bacterial infection. Complete demographic and clinical metadata are available
3. 4- Enough urine volume for molecular assessment.

### Exclusion Criteria

1. Exclusion criteria included any of the following:
2. The metadata is missing or the clinical record is incomplete.
3. To low urine quantity and late urine collection for DNA extraction and sequencing.
4. Sample contamination or degraded state of the genomic DNA.

### Collection of clinical and demographic data

Demographic and clinical data, including age, sex, BMI, diabetes status, and urine culture results as well as antibiotic usage over the past seven days were collected from patient records and documented for each sample. These variables were used to describe baseline cohort characteristics and defined clinically relevant comparison groups for microbiome analysis.

Continuous variables (age and BMI) were described as mean with standard deviation, whereas categorical variables (sex, diabetes status, culture results and exposure to antibiotics) were expressed in frequency with percentages. This where the principle of reporting permits general overview representation about clinical context of sequenced UTI samples.

### Urine Sample Collection and Handling

Samples were obtained in aseptic conditions (midstream urine) with the use of sterile containers to avoid external contamination. Specimens were quickly transferred to the laboratory and underwent processing under controlled conditions. Aliquots were set aside for routine microbiological culture and molecular analysis. All samples were stored at suitable temperatures prior to DNA extraction in order to maintain the microbial integrity.

### DNA Extraction and 16S rRNA Gene Sequencing

Total microbial DNA was extracted from urine samples using a standardized genomic DNA extraction protocol that has been optimized for low-biomass clinical specimens. DNA extraction was quantified and evaluated for purity before further processing.

Using primers that target conserved hypervariable regions ideal for genus-level taxonomic profiling, we amplified the bacterial 16S rRNA gene. The amplified products underwent next-generation sequencing to obtain high-throughput reads that reflect each urine sample’s bacteria composition. The sequencing workflow allowed culture-free identification of microbial taxa and the quantification of their relative abundance in every sample.(36)

### Bioinformatics Processing and Taxonomic Assignment

Low quality reads and possible artifacts were filtered out from raw sequencing data. Chimeric sequences were removed to allow high-resolution taxonomic assignment. Filtered reads were subsequently clustered into operational taxonomic units (OTUs) or amplicon sequence variants (ASVs) based on sequence similarity thresholds.

Taxonomic assignment of bacterial sequences at the genus level was conducted using curated reference databases. Summary tables for relative abundance were generated for all categories, enabling dominant genera identification and detailed profiling of microbe composition within samples.(37)

### Microbiome Composition Analysis

#### Dominant Genus Identification

The dominant genus was defined as the bacterial genus with the dominant relative abundance in that sample. The dominance proportion was computed to determine the strength of taxon driven community structure. Distinct values of dominance showed degree of evenness and strong pathogen predominance (higher) as opposed to mixed microbial communities (lower).

#### Within-Sample Relative Abundance Profiles

Top bacterial genera in each sample were determined from ranked relative abundance values. These rank-abundance profiles were utilized to delineate bacterial community structure steepness and for detection of variant taxa present in all the patients, as a measure of inter-individual heterogeneity of urinary microbiota.

#### Cohort-Level Genus Summary

To obtain cohort-wide microbial signatures, we calculated the mean relative abundance and prevalence of each genus in all samples. Mean relative abundance provided the average contribution of every genera on the overall community whereas prevalence indicated the proportion of samples in which that specific genus was detected. Individual or metagenomic abundance measurement and additional analysis of these taxa as described above, provided identification of genera that were consistently present or selectively enriched in subsets of patients.(38)

### Alpha Diversity Analysis

Alpha diversity was assessed by calculating within-sample microbial diversity indices. The Shannon index (H) was used to quantify both richness and evenness, and the Simpson index allowed an additional assessment of dominance structure. Observed species counts were defined as the number of genera detected in each environmental sample, and a Chao1 based richness estimator was used to estimate unseen richness from rare taxa. Metrics were employed to classify whether UTI samples demonstrated single-taxon dominance or more complex polymicrobial compositions.

### Beta Diversity and Ordination Analysis

We used the Bray–Curtis distance metric at the genus-level to determine between-sample compositional differences. Pairwise distance matrices were calculated to measure the similarity or differences in microbial community composition among samples with values close to zero indicating greater similarities and closer to one suggesting significant compositional dissimilarities.

Low-dimensional ordination plots were obtained via Principal Coordinates Analysis (PCoA) using the Bray–Curtis distance matrix. This strategy allowed visualization and exploration of clustering patterns between samples and relationships between microbial profiles and clinical metadata.

## Statistical Analysis

Performed descriptive statistical analysis describing demographic, clinical and microbiome variables. Continuous variables were presented as mean ± standard deviation, while categorical variables were reported as frequencies (n) and percentages (%). Qualitative measures of diversity, such as diversity indices and relative abundances were analyzed using exploratory statistical methods to identify variability between samples.

**Table 1.**
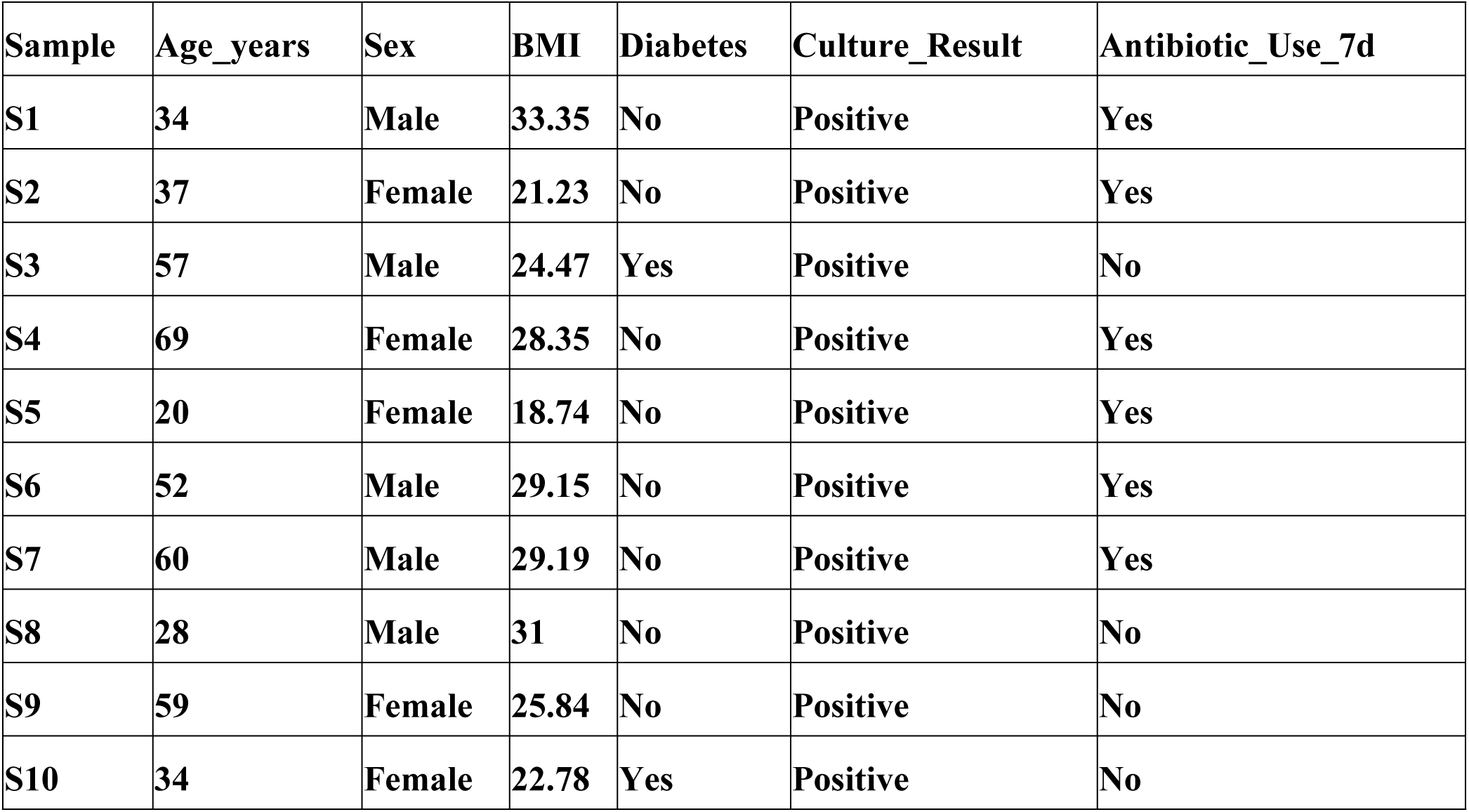
Demographic and Clinical Characteristics by Sample.

## Results

The following table presents the sample-associated is a demographic and clinical metadata schema of 10 sequenced samples (S1-S10). These variables must be defined from the patient records and used in a publishable analysis in two ways (i) description of the baseline cohort (distribution of age, balance of sex, profile of comorbidity, positivity for cultures and previous antibiotic exposure), and (ii) specification of clinically important comparison groups36(i.e., culture-positive vs culture-negative; other severe vs mild symptoms; exposed to antibiotics pre-illness vs non-exposed). Having exact sample IDs is critical as it links metadata and 16S outputs

**Table 2.**
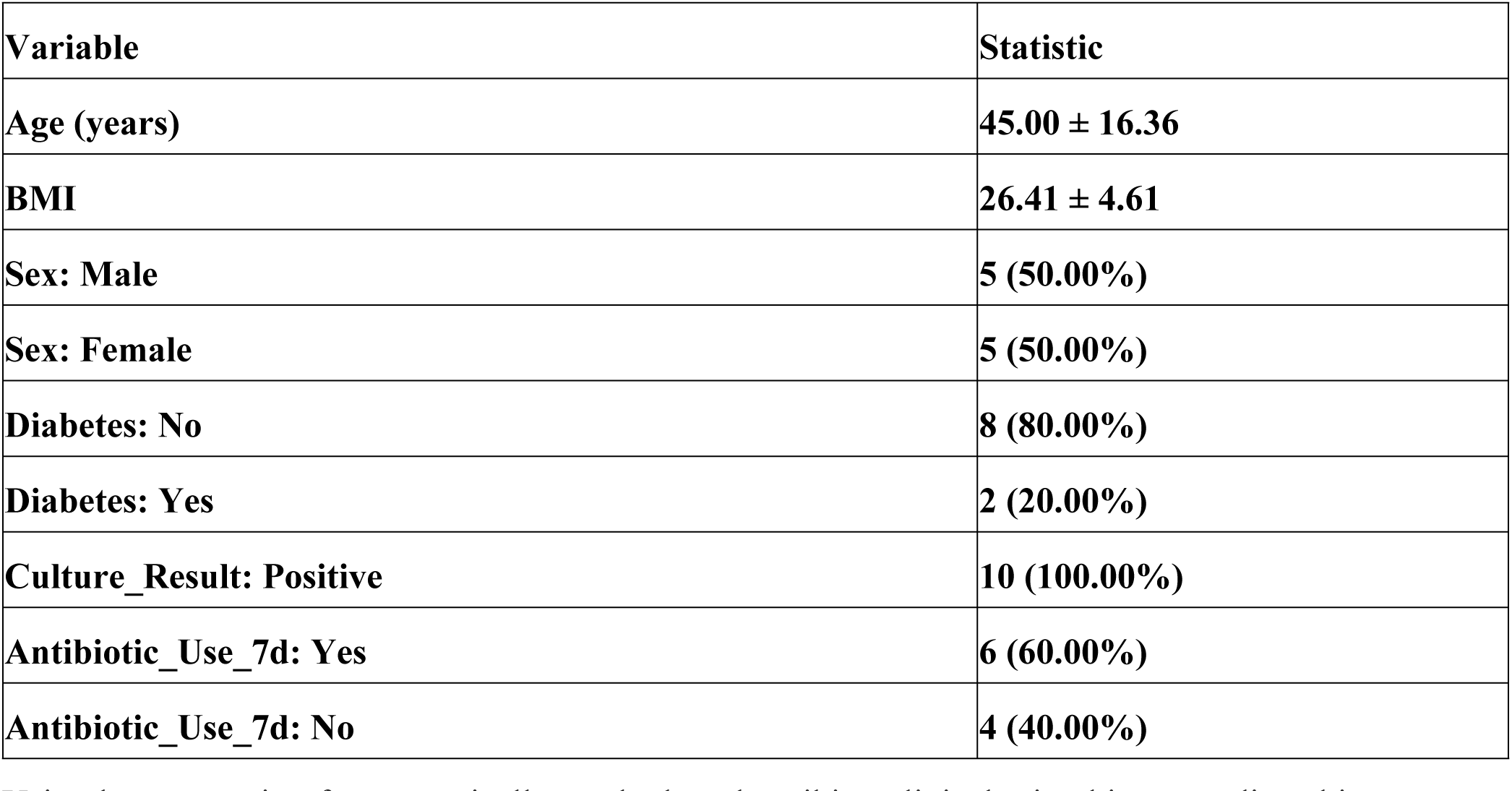
Demographic Summary (Mean±SD and n (%)

Using base reporting format typically used when describing clinical microbiome studies, this summary table will report continuous variables as mean standard deviation and categorical variables as n (%) After the real metadata are properly positioned, this table is the main output of cohort-give an account of that will help readers to understand the clinical background of sequenced test example and for deciding on, whether principle factors are adjusted in any one examination gatherings.

**Table 3.** Dominant Bacterial Genus in Each Sample and Relative Abundance.

| **Sample** | **Dominant Genus** | **Dominant_RelAbun** |
| --- | --- | --- |
| **S1** | **Klebsiella** | **0.86** |
| **S2** | **Pseudomonas** | **0.86** |
| **S3** | **Klebsiella** | **0.49** |
| **S4** | **Proteus** | **0.46** |
| **S5** | **Escherichia** | **0.68** |
| **S6** | **Pseudomonas** | **1.00** |
| **S7** | **Pseudomonas** | **1** |
| **S8** | **Pseudomonas** | **1.00** |
| **S9** | **Proteus** | **0.68** |
| **S10** | **Burkholderia-Burkholderia-Paraburkholderia** | **0.68** |

Relative abundance is indicated and the predominant genus in each sample is determined. The controlling genus directly summarizes diagnostics at the genus level, and dominance ratio is a measure of strength by which single genus describes sample profile. Higher dominance values indicate a more consolidated community structure in one taxon (lower evenness), and lower dominance values indicate that many genera still have function to provide. The table is particularly useful to report samples likely uropathogens, and to screen mixed and/or atypical communities.

**Table 4.** Top 10 Bacterial Genera per Sample (Relative Abundance Profile)

| **Genus** | **Sample** | **RelAbun** |
| --- | --- | --- |
| **Klebsiella** | **S1** | **0.86** |
| **Escherichia** | **S1** | **0.10** |
| **Enterococcus** | **S1** | **0.04** |
| **Rhodococcus** | **S1** | **0.00** |
| **Synechococcus** | **S1** | **0** |
| **Brevibacillus** | **S1** | **0** |
| **Paenibacillus** | **S1** | **0** |
| **Staphylococcus** | **S1** | **0** |
| **Lactobacillus** | **S1** | **0** |
| **Clostridium** | **S1** | **0** |
| **Burkholderia-Burkholderia-Paraburkholderia** | **S10** | **0.68** |
| **Enterococcus** | **S10** | **0.29** |
| **Burkholderia-Paraburkholderia** | **S10** | **0.03** |
| **Pseudomonas** | **S10** | **0.00** |
| **Rhodococcus** | **S10** | **0** |
| **Synechococcus** | **S10** | **0** |
| **Brevibacillus** | **S10** | **0** |
| **Paenibacillus** | **S10** | **0** |
| **Staphylococcus** | **S10** | **0** |
| **Lactobacillus** | **S10** | **0** |
| **Pseudomonas** | **S2** | **0.86** |
| **Brevundimonas** | **S2** | **0.13** |
| **Paenibacillus** | **S2** | **0.01** |
| **Ochrobactrum** | **S2** | **0.00** |
| **Rhodococcus** | **S2** | **0** |
| **Synechococcus** | **S2** | **0** |
| **Brevibacillus** | **S2** | **0** |
| **Staphylococcus** | **S2** | **0** |
| **Enterococcus** | **S2** | **0** |
| **Klebsiella** | **S3** | **0.49** |
| **Lactobacillus** | **S3** | **0.29** |
| **Escherichia** | **S3** | **0.17** |
| **Enterobacter** | **S3** | **0.04** |
| **Enterococcus** | **S3** | **0.00** |
| **Pseudomonas** | **S3** | **0.00** |
| **Rhodococcus** | **S3** | **0** |
| **Synechococcus** | **S3** | **0** |
| **Brevibacillus** | **S3** | **0** |
| **Paenibacillus** | **S3** | **0** |
| **Proteus** | **S4** | **0.46** |
| **Pseudomonas** | **S4** | **0.34** |
| **Enterococcus** | **S4** | **0.20** |
| **Lactobacillus** | **S4** | **0.00** |
| **Thalassospira** | **S4** | **0.00** |
| **Rhodococcus** | **S4** | **0** |
| **Synechococcus** | **S4** | **0** |
| **Brevibacillus** | **S4** | **0** |
| **Paenibacillus** | **S4** | **0** |
| **Staphylococcus** | **S4** | **0** |
| **Escherichia** | **S5** | **0.68** |
| **Enterococcus** | **S5** | **0.25** |
| **Pseudomonas** | **S5** | **0.04** |
| **Klebsiella** | **S5** | **0.02** |
| **Stenotrophomonas** | **S5** | **0.00** |
| **Brevibacillus** | **S5** | **0.00** |
| **Staphylococcus** | **S5** | **0.00** |
| **Rhodococcus** | **S5** | **0** |
| **Synechococcus** | **S5** | **0** |
| **Paenibacillus** | **S5** | **0** |
| **Pseudomonas** | **S6** | **1.00** |
| **Brevundimonas** | **S6** | **0.00** |
| **Rhodococcus** | **S6** | **0** |
| **Synechococcus** | **S6** | **0** |
| **Brevibacillus** | **S6** | **0** |
| **Paenibacillus** | **S6** | **0** |
| **Staphylococcus** | **S6** | **0** |
| **Enterococcus** | **S6** | **0** |
| **Lactobacillus** | **S6** | **0** |
| **Clostridium** | **S6** | **0** |
| **Pseudomonas** | **S7** | **1** |
| **Rhodococcus** | **S7** | **0** |
| **Synechococcus** | **S7** | **0** |
| **Brevibacillus** | **S7** | **0** |
| **Paenibacillus** | **S7** | **0** |
| **Staphylococcus** | **S7** | **0** |
| **Enterococcus** | **S7** | **0** |
| **Lactobacillus** | **S7** | **0** |
| **Clostridium** | **S7** | **0** |
| **Brevundimonas** | **S7** | **0** |
| **Pseudomonas** | **S8** | **1.00** |
| **Rhodococcus** | **S8** | **0** |
| **Synechococcus** | **S8** | **0** |
| **Brevibacillus** | **S8** | **0** |
| **Paenibacillus** | **S8** | **0** |
| **Staphylococcus** | **S8** | **0** |
| **Enterococcus** | **S8** | **0** |
| **Lactobacillus** | **S8** | **0** |
| **Clostridium** | **S8** | **0** |
| **Proteus** | **S9** | **0.68** |
| **Enterococcus** | **S9** | **0.32** |
| **Pseudomonas** | **S9** | **0.00** |
| **Clostridium** | **S9** | **0.00** |
| **Synechococcus** | **S9** | **0.00** |
| **Rhodococcus** | **S9** | **0** |
| **Brevibacillus** | **S9** | **0** |
| **Paenibacillus** | **S9** | **0** |
| **Staphylococcus** | **S9** | **0** |
| **Lactobacillus** | **S9** | **0** |

This table contains the most prevalent genera for each sample, as well as within sample composition. This profile can be demonstrated with a rank-abundance pattern that indicates if it is steep (i.e., few genera are dominant with many low-abundance taxa) or more gradual which is deemed as higher evenness. By comparing these profiles in the samples, inter-patient heterogeneity can be evaluated via a descriptive approach and identify genera shared by several patients. From the top-10 list, sulci also provide a short list of potent targets for evidence-based reporting and downstream between-group testing once clinical groups are defined.

**Table 5.** Top Bacterial Genera Across All Samples (Mean Relative Abundance and Prevalence)

| **Genus** | **Mean_RelAbun** | **Prevalence** |
| --- | --- | --- |
| **Pseudomonas** | **0.42** | **90** |
| **Klebsiella** | **0.14** | **30** |
| **Proteus** | **0.11** | **20** |
| **Enterococcus** | **0.11** | **60** |
| **Escherichia** | **0.10** | **30** |
| **Burkholderia-Burkholderia-Paraburkholderia** | **0.07** | **10** |
| **Lactobacillus** | **0.03** | **20** |
| **Brevundimonas** | **0.01** | **20** |
| **Enterobacter** | **0.00** | **10** |
| **Burkholderia-Paraburkholderia** | **0.00** | **10** |
| **Paenibacillus** | **0.00** | **10** |
| **Rhodococcus** | **0.00** | **10** |
| **Stenotrophomonas** | **0.00** | **10** |
| **Brevibacillus** | **0.00** | **10** |
| **Clostridium** | **0.00** | **10** |
| **Ochrobactrum** | **0.00** | **10** |
| **Staphylococcus** | **0.00** | **10** |
| **Thalassospira** | **0.00** | **10** |
| **Synechococcus** | **0.00** | **10** |

This table gives summary cohort-level genus patterns across two complementary statistics; mean relative abundance and prevalence. Mean relative abundance is the arithmetic mean of signal strengths in the cohort, and prevalence indicates how consistently a genus was detected in the samples. Genera with both high mean abundance and high prevalence represent strong cohort-wide signals, whereas genera that are highly abundant but not prevalent indicate the enrichment of a subgroup of patients. Reporting the 2 metrics in conjunction with each other has its benefits and these are significantly better chronology of the data being analysed alongside an objective explanation for why taxa were chosen, specifically within metadata based groups.

**Table 6.** Alpha Diversity Indices Across Samples (Shannon, Simpson, Observed Features, Chao1)

| Sample | shannon | simpson | observed_species | chao1 |
| --- | --- | --- | --- | --- |
| S1 | 3.66 | 0.87 | 36 | 36 |
| S2 | 1.08 | 0.30 | 25 | 25 |
| S3 | 4.64 | 0.94 | 50 | 50 |
| S4 | 2.39 | 0.76 | 12 | 12 |
| S5 | 3.01 | 0.85 | 18 | 18 |
| S6 | 1.52 | 0.63 | 4 | 4 |
| S7 | 0.65 | 0.28 | 3 | 3 |
| S8 | 1.27 | 0.48 | 8 | 8 |
| S9 | 3.41 | 0.87 | 19 | 19 |
| S10 | 3.66 | 0.87 | 22 | 22 |

Here the indices of within-sample (alpha) diversity reflect genus-level profiles richness and evenness. It is called Shannon because Shannon grows in both evenness and richness meaning that a lower value of varies in genera dominance will result into few genera dominating the presence while higher values of sequence data which gives high varaibility between serotype as well as type to dominate print. With Shannon filling in a void on the dominance structure, Simpson has more or less exactly filled that in. Observed features is a measure of determined richness and chao1 is an estimate of undetermined richness based on rare taxa When real metadata is provided, they can be statistically compared between clinical groups, and allow for descriptive statements to be made (for example whether UTI samples tend to consist of the dominance of single taxa or mixed communities).

**Table 7.** Bray–Curtis Distance Matrix (Genus Level)

| Sample | S1 | S2 | S3 | S4 | S5 | S6 | S7 | S8 | S9 | S10 |
| --- | --- | --- | --- | --- | --- | --- | --- | --- | --- | --- |
| S1 | 0 | 1 | 0.40 | 0.96 | 0.84 | 1 | 1 | 1 | 0.96 | 0.96 |
| **S2** | **1** | **0** | **1.00** | **0.66** | **0.96** | **0.14** | **0.14** | **0.14** | **1.00** | **1.00** |
| **S3** | **0.40** | **1.00** | **0** | **0.99** | **0.80** | **1.00** | **1.00** | **1.00** | **0.99** | **0.99** |
| **S4** | **0.96** | **0.66** | **0.99** | **0** | **0.76** | **0.66** | **0.66** | **0.66** | **0.34** | **0.80** |
| **S5** | **0.84** | **0.96** | **0.80** | **0.76** | **0** | **0.96** | **0.96** | **0.96** | **0.74** | **0.74** |
| **S6** | **1** | **0.14** | **1.00** | **0.66** | **0.96** | **0** | **0.00** | **0.00** | **1.00** | **1.00** |
| **S7** | **1** | **0.14** | **1.00** | **0.66** | **0.96** | **0.00** | **0** | **0.00** | **1.00** | **1.00** |
| **S8** | **1** | **0.14** | **1.00** | **0.66** | **0.96** | **0.00** | **0.00** | **0** | **1.00** | **1.00** |
| **S9** | **0.96** | **1.00** | **0.99** | **0.34** | **0.74** | **1.00** | **1.00** | **1.00** | **0** | **0.71** |
| **S10** | **0.96** | **1.00** | **0.99** | **0.80** | **0.74** | **1.00** | **1.00** | **1.00** | **0.71** | **0** |

This table provides pairwise BrayCurtis distances of samples, i.e. between sample compositional similarity at the genus level (0 = identical; 1 = totally different). For shorter distances (small samples) these results suggest those samples shared similar dominant and subordinate genera while the longer distances (large shift) indicate that significant compositional changes took place in the involved samples. Once added, both ordination (PCOA) and inferential testing (e.g., PERMANOVA) are contingent to the distance matrix.

**Table 8.** PCoA Coordinates Based on Bray–Curtis Dissimilarity (Genus Level)

| **Sample** | **PC1** | **PC2** |
| --- | --- | --- |
| **S1** | **-0.41** | **0.47** |
| **S2** | **0.46** | **0.06** |
| **S3** | **-0.40** | **0.51** |
| **S4** | **0.03** | **-0.35** |
| **S5** | **-0.38** | **-0.09** |
| **S6** | **0.46** | **0.06** |
| **S7** | **0.46** | **0.06** |
| **S8** | **0.46** | **0.06** |
| **S9** | **-0.34** | **-0.45** |
| **S10** | **-0.35** | **-0.32** |

The table shows the two initial PCoA values achieved in Bray-Curtis distances. PCoA provides a low-dimensional embedding of the differences between samples: Samples that are closer to each other in this space show greater similarity with respect to genus profiles while more distally located (more dissimilar) samples indicate a higher degree of compositional dissimilarity. With real clinical meta data, we can formalize the ordination to visually examine group clustering, while justifying testing with PERMANOVA for the extent of group separation.

**Figure 1.**
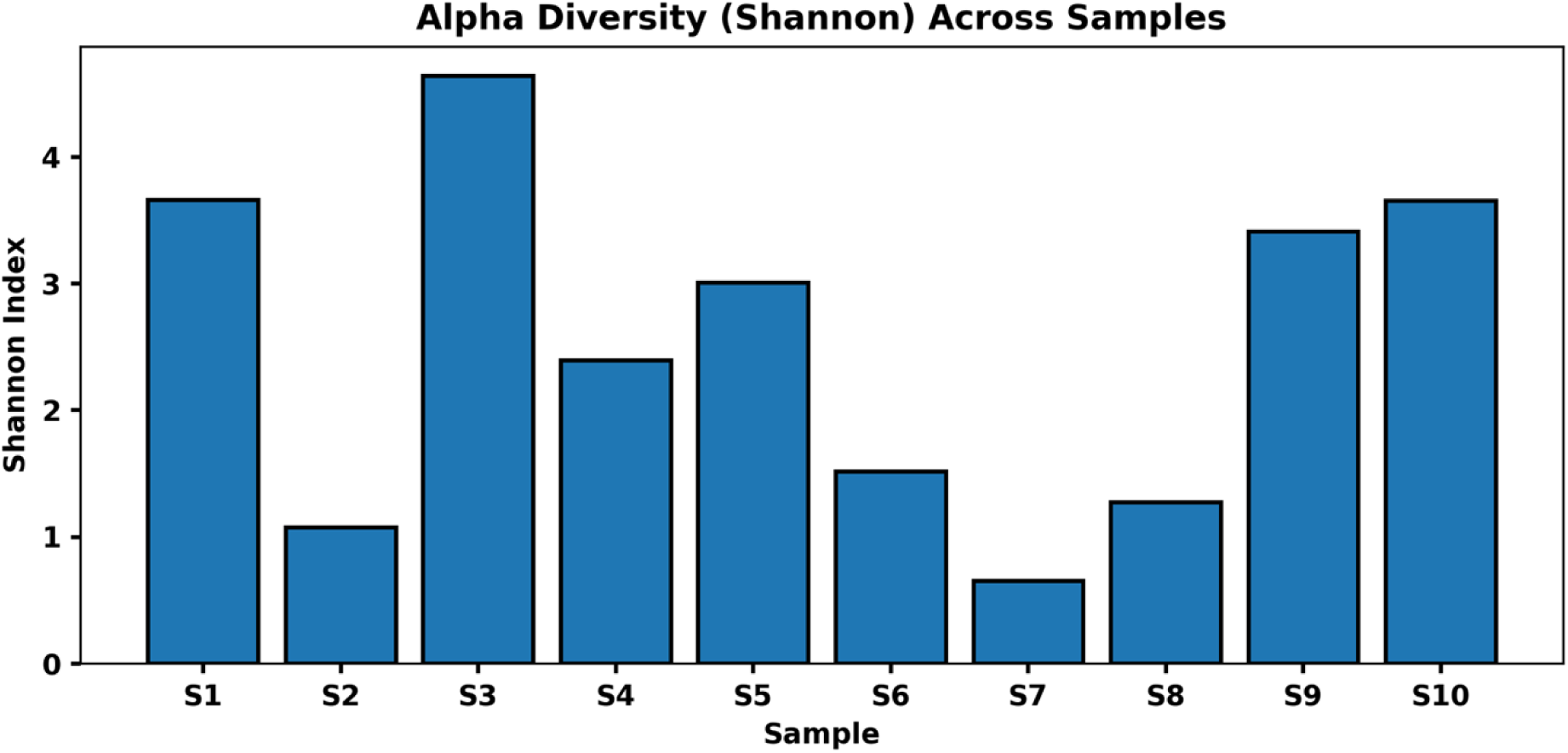
Alpha Diversity Across Samples (Shannon Index) This figure shows the Shannon alpha diversity index across the 10 samples. Higher Shannon values suggest higher within-sample diversity and evenness; lower values imply dominance by fewer genera. The figure is amenable to visual comparison for rapid assessment across patients and will also allow for between-group comparisons after clinical grouping variables are ascertained.

## Discussion

Here we present the first pilot study of urinary microbiome composition and diversity in clinician confirmed urinary tract infection (UTI) samples using a high-throughput 16S rRNA sequencing approach. Through combining demographic metadata, genus-level taxonomic profiles, and ecological diversity metrics, the work provides valuable insights into the heterogeneity of microbial communities associated with UTIs and the realization that culture-independent microbiome analysis can yield important benefits.

The cohort demographic and clinical characteristics showed sex balance, mean age of 45 years and body mass index moderately dispersed. The majority of the participants were non-diabetic and all cases were culture-positive, confirming active infection at the time of sampling. Importantly, recent exposure to antibiotic medication was reported in 60% of patients (an important consideration for interpreting patterns microbiome composition and diversity)(22). It is well-known that antibiotic treatment modifies microbial ecology (while reducing susceptible taxa it facilitates the overgrowth on resistant or opportunistic organisms)(23) what may contribute to dominance patterns documented in some samples.

One of the principal results of this study is the remarkable inter-individual variability in dominant bacterial genera observed in UTI samples. Overall, Pseudomonas was the most prevalent identified genus and this contributed with particularly high relative abundance values in several samples indicating community dominance, while other classical uropathogenic bacteria (e.g. Klebsiella, Proteus and Escherichia) predominated among some isolated patients. Such a pattern shows that UTIs are not microbiologically homogeneous entities; they represent a continuum of infections with discrete microbial compositions which vary from one patient to the next. These findings of Burkholderia-related taxa being the dominant genus in one such sample also lend support to the concept that atypical or less commonly recognized genera may contribute to infection in some clinical contexts(24).

within-sample composition analyses revealed steep rank-abundance distributions in many if not most (one genus capturing the bulk of the microbial signal with surrounding taxa at very low relative abundances). All of these profiles show the hallmarks of a pathogen-driven low evenness infection consistent with classic acute UTI pathogenesis, in which a single uropathogen colonizes and inhibits other competing microorganisms(25,26). However, some samples evidenced more gradual rank-abundance curves with detectable secondary taxa (e.g. Enterococcus or Lactobacillus) indicative of polymicrobial communities consistent with either co-infection, transitional dysbiosis, and/or persistence of commensal flora despite pathogen predominance.

Overall, cohort-level genus summaries supported the presence of predominant but repeatable microbial signatures across patients. Pseudomonas had not only the highest mean relative abundance, but it was also the most prevalent which indicates that it is a consistent and strong presence within the cohort. In contrast, genera including Klebsiella, Proteus and Escherichia were moderately prevalent with variable abundance indicating that these organisms would be relatively dominant in some closely related individuals but absent from others.(27) Relatively high prevalence yet moderate mean abundance placed Enterococcus as a recurrent but not necessarily dominant taxon in UTI microbiomes. This pattern of mean abundance and prevalence gives a more nuanced account of cohort-wide microbial patterns compared to the reliance on dominance(28).

Alpha diversity evaluation showed significant heterogeneity of richness and evenness among samples. Some samples showed very low Shannon and Simpson index values, indicating near-complete dominance by a single genus, while others had higher diversity indices and greater numbers of observed features(29). These results illustrate that UTIs are linked to low-diversity pathogen-dominated communities, as well as higher-diversity polymicrobial ecosystems. Notably, the detection of mixed communities in multiple samples raises questions about the traditional view of UTIs as strictly mono-microbial infections and indicates that interactions between microbial species within the urinary tract could contribute to disease progression, persistence or recurrence(30).

Beta diversity analysis based on Bray–Curtis distances showed significant compositional differences between many samples with multiple pairwise distances reaching close to their maximum values. Thus, it shows that the microbial community structures were different from each other between patients, supports “individualized microbiome signatures” in UTI. In contrast, some very small distances between specific samples, in particular dominated by Pseudomonas suggested that infections associated with the same dominant genus may result in more similar community arrangements. The results were also mirrored in PCoA ordination results, where clustering of specific samples suggested shared dominant taxa and dispersion of others reflected unique compositional profiles(31).

Also, from a clinical perspective, the heterogeneity of urinary microbiome composition suggests that different microbiomes may have important diagnostic and therapeutic considerations. dominance of classical uropathogens in a number of samples supports the ongoing use of targeted antimicrobial therapy. Yet, the identification of mixed microbial communities and atypical taxa could also indicate limitations of traditional culture methods that may fail to capture the entire microbial landscape(4). This implies that molecular microbiome profiling may provide additional insights beyond conventional diagnostics from co-occurring organisms that could alter treatment response or recurrence risk.

Also interesting, recent antibiotic exposure may affect a subjects pattern of dominance/diversity. Dominance of single genera was often strong in samples from antibiotic-exposed patients, reflecting selective pressure for resistant or opportunistic organisms. Antibiotic-naïve samples, however, showed greater diversity and more even community structures. While the small sample size prevents definitive conclusions, these trends indicate that antibiotic history may represent an important covariate in urinary microbiome studies, as well as a possible explanation for inter-patient variability(32,33).

These findings from this pilot study should be interpreted considering the following limitations. The somewhat small sample size limits statistical power and prevents strong subgroup comparisons by clinical variables (e.g. diabetes status, prior antibiotic exposure)(34). Furhtermore, the cross-sectional approach records microbiome composition at a single time point and does not reveal temporal dynamics or longitudinal changes throughout therapy/recurrence. Moreover, significant inflations of functional richness have been reported in some studies and genus-level resolution that characterizes 16S rRNA sequencing does not provide sufficient detail for identifying functions at the species level or describing taxa that were identified and their function(35).

Introducing phenomena such as these may limit the extent to which this study can realistically be extrapolated to generalized UTI patients but nevertheless provides useful preliminary evidence that urinary microbiomes in individuals with UTI are diverse and many dominated by strong pathogen presence intermingled amidst polymicrobial community structures(30). Dominance analysis, diversity indices and compositional dissimilarity metrics integrate data into an ecophysiological perspective of UTI-associated microbiota. These results corroborate the notion that UTIs are a microbiologically heterogeneous disease that is influenced by host aetiology/underlying factors and previous antimicrobial use rather than identical infection caused by one pathogen across all patients(8).

Larger cohorts and longitudinal sampling are needed, but further studies in this area may lead to larger breakthroughs exploring associations between microbiome patterns and clinical phenomenology such as symptom severity, treatment response, and recurrence risk. Inclusion of granular clinical metadata and sophisticated multi-omics methods may further uncover the functional role of urinary microbiota in UTI pathogenesis, ultimately aiding in the development of microbiome-informed diagnostic and therapeutic strategies.

## Ethical Considerations

Confidentiality was maintained by anonymizing all patient data and coding with unique sample identifiers. The study used de-identified clinical specimens that were originally obtained as part of routine diagnostics, and all analyses were performed in accordance with the institutional and ethical guidelines for human-derived clinical material.

## Conclusion

This pilot study has characterized urine microbiome communities and compositions in clinically diagnosed urinary tract infection (UTI) samples with 16S rRNA gene sequencing. These results show considerable patient variability in UTI-associated microbial communities with respect to dominant genera, diversity indices, and general community structure. There was great heterogeneity among samples from different individuals with respect to making up the dominant taxa, yet classical uropathogens (e.g., Pseudomonas, Klebsiella, Proteus, Escherichia) can be found as dominant in various individuals. The presence of secondary, low-abundance, or rare genera suggests that more complex microbial ecosystems can be present at UTIs than would be identified by standard cultures alone.

## Funding

This study funded by ZYMO RESEARCH CORP. U.S.A, SMRT Grant project, in00650

## Acknowledgment

We would like to thank ZYMO RESEARCH CORP., U.S.A. for their support in conducting the 16S rRNA analysis for this study. Their expertise and resources were instrumental in achieving the results presented in this manuscript.

## Authors’ Contributions

MusaabHamid Farhan, Ayad M.J. Al-Mamoori, and Noor S.K. Al-Khafaji designed the study. Musaab Hamid Farhan and Aws H. J. Al_Rahhal collected and analyzed the data. Musaab Hamid Farhan and Ayad M.J. Al-Mamoori interpreted the results. Musaab Hamid Farhan drafted the manuscript. All authors reviewed, edited, and approved the final manuscript.

## Notes

### Competing Interest Statement

The authors have declared no competing interest.

